# Epigenetic regulation underlying *Plasmodium berghei* gene expression during its developmental transition from host to vector

**DOI:** 10.1101/646430

**Authors:** Kathrin Witmer, Sabine AK Fraschka, Dina Vlachou, Richárd Bártfai, George K Christophides

## Abstract

Epigenetic regulation of gene expression is an important attribute in the survival and adaptation of the malaria parasite *Plasmodium* in its human host. Our understanding of epigenetic regulation of gene expression in *Plasmodium* developmental stages beyond asexual replication in the mammalian host is sparse. We used chromatin immune-precipitation (ChIP) and RNA sequencing to create an epigenetic and transcriptomic map of the murine parasite *Plasmodium berghei* development from asexual blood stages to male and female gametocytes, and finally, to ookinetes. We show that heterochromatin 1 (HP1) almost exclusively associates with variantly expressed gene families at subtelomeric regions and remains stable across stages and various parasite lines. Variant expression based on heterochromatic silencing is observed only in very few genes. In contrast, the active histone mark histone 3 Lysine 9 acetylation (H3K9ac) is found between heterochromatin boundaries and occurs as a sharp peak around the start codon for ribosomal protein genes. H3K9ac occupancy positively correlates with gene transcripts in asexual blood stages, male gametocytes and ookinetes. Interestingly, H3K9ac occupancy does not correlate with transcript abundance in female gametocytes. Finally, we identify novel DNA motifs upstream of ookinete-specific genes thought to be involved in transcriptional activation upon fertilization.

## INTRODUCTION

Malaria is caused by apicomplexan parasites of the genus *Plasmodium* and is transmitted to humans through bites of anopheline mosquitoes. Malaria clinical cases and deaths decreased significantly over the past decade but began to stabilize since 2015 indicating that current measures have now reached their maximum capacity and that new measures are urgently needed (1). Parasite transmission through the mosquito vector is a natural bottleneck in its development and a favourable stage for interventions aiming at malaria control and elimination. Therefore, research towards understanding parasite development in the mosquito has been intensified in recent years.

Haploid parasites cyclically infect and asexually replicate in the infected red blood cells (iRBCs) of the host causing disease pathology. Eventually, some parasites escape this cycle and form sexual forms called gametocytes, the stage infective to mosquitoes. In the mosquito gut lumen, ingested gametocytes are activated to form gametes. Female gametocytes exit iRBCs, and messenger RNAs (mRNAs), stored in a messenger ribonucleoprotein (mRNP) complex, become available for translation (2, 3). Activation of male gametocytes involves three rapid rounds of endomitosis leading to eight flagellated microgametes, a process called exflagellation (4). After fertilization, the zygote embarks on a meiotic endoreplication cycle while traversing the mosquito midgut epithelium in the form of an ookinete that upon arrival at the midgut basal side transforms into an oocyst (5). Over the next days, endomitotic replication in the oocyst produces hundreds of sporozoites that, upon oocyst rupture, travel to the mosquito salivary gland for inoculation into a vertebrate host with the next mosquito bite.

Epigenetic regulation is important for parasite survival within the human host (6). Genes involved in host-parasite interactions or coding for virulence factors or ligands involved in RBC invasion are epigenetically regulated (7, 8), while some genes involved in drug resistance are epigenetically switched on or off in an environment-dependent manner (9). Importantly, expression of parasite antigens in the mouse host is reset after its passage through the mosquito, suggesting that epigenetic imprinting may be erased during the mosquito stage (10).

*P. falciparum* HP1 (*Pf*HP1) is shown to bind exclusively to histone 3 lysine 9 tri-methylation (H3K9me3) and be the hallmark of transcriptionally silent heterochromatin (7, 11). Heterochromatic loci are largely confined to telomeric and subtelomeric regions as well as chromosome-central islands and almost invariably associated with variantly expressed multigene protein families (7, 11–15). In contrast, the universal histone mark linked to euchromatin H3K9ac is shown to be related to active transcription in *P. falciparum* asexual blood stages (12, 14).

The binding of transcription factors on the gene promoter region is heavily dependent on the state of the surrounding chromatin. Members of the apicomplexan-specific ApiAP2 family of transcription factors are found to control major cell fate decision events in the parasite lifecycle, in addition to housekeeping processes (16–21). AP2-G is shown to be the master regulator of gametocytogenesis (20, 21), activating a number of gametocyte-specific genes (22). Similarly, the ookinete-specific AP2-O, which is itself regulated by the mRNP complex, activates transcription of over 400 genes needed for ookinete development and mosquito midgut traversal (17, 23). Three additional ookinete-specific ApiAP2 transcription factors have been identified, which play a role just before or after ookinete formation (16).

Here, we investigate how epigenetic traits change in the murine malaria parasite *P. berghei* during its transition from the murine host to the mosquito vector, using asexual blood stages (ABS), female (FG) and male (MG) gametocytes, and ookinetes (OOK). We find a conserved distribution of heterochromatin through parasite development, and surprisingly little difference between different parasite lines. In all stages, heterochromatin is confined to subtelomeric regions, except two chromosomal central loci encompassing genes encoding the oocyst capsule protein Cap380 and a conserved protein of unknown function (PbANKA_0934600). Implementing transcriptomics we identify that variant expression of multigene families genes occurs in all developmental stages and not only in ABS. In stark contrast to heterochromatin, the distribution of H3K9ac is dynamic throughout parasite development, and peaks around the start codon of ribosomal protein genes. In consistence with previous findings, H3K9ac enrichment in gene 5’ untranslated regions (5’UTRs) correlates with transcript levels in ABS. Additionally, H3K9ac enrichment correlates with increased transcript abundance in MG and OOK but not in FG suggesting a different epigenetic state of FG compared to other developmental stages. Finally, we identify four novel DNA motifs in addition to the AP2-O enriched in the 5’UTR of OOK-specific genes, suggesting that additional transcription factors are involved in orchestrating transcription in OOK. Our study adds substantially to our understanding of malaria transmission biology.

## MATERIAL AND METHODS

### Parasites

*P. berghei* clone *ANKA 2.33* (for asexual blood stages) (24), the *507m6cl1 (c507)* line (for ookinetes) (25) were maintained in 6–8 week old female Tuck’s Ordinary (TO) (Harlan, UK). The *820cl1m1cl1 (wt-fluo)* line (for gametocytes) (26) was maintained in 6–8 week old female CD1 mice (Harlan, UK).

### Chromatin extraction and fragmentation

Asexual blood stage parasites were harvested via heart puncture and passed through a Plasmodipur filter (Europroxima) to remove leucocytes, resuspended in RPMI-1640 medium (Sigma-Aldrich) and crosslinked with 1% formaldehyde in PBS for 10min at 37°C. Crosslinking was quenched adding glycine to an end concentration of 0.125M. RBC were then lysed with 0.15% saponin (in PBS) for 5-10 minutes on ice. To obtain nuclei the resulting parasite pellet was lysed with cell lysis buffer (20 mM Hepes, 10mM KCl,1mM EDTA, 1mM EGTA, 0.65% NP-40, 1mM DTT, 1x protease inhibitor (Roche)). The pellet was resuspended in sonication buffer (1% SDS, 50mM Tris pH8, 10 mM EDTA, 1x protease inhibitor (cOmplete™, Mini, EDTA-free, Roche)) and sheared for 25 minutes (30sec ON, 30 sec OFF; settings HIGH) using a Bioruptor® Plus sonication device (Diagenode) to obtain DNA fragments of around 100-300bp.

The method for the purification of gametocytes was modified from (27). Briefly, mice were pretreated by intra peritoneal injection of 0.2 ml phenylhydrazine (6 mg/ml in PBS) to stimulate reticulocyte formation two days prior to infection with parasites. Gametocyte-enriched blood was harvested via heart puncture and blood was immediately resuspended in 4°C coelenterazine loading buffer (CLB) (1x PBS, 20 mM HEPES, 20 mM glucose, 4 mM sodium bicarbonate, 1 mM EDTA, 0.1% BSA in PBS, pH 7.24–7.31) and magnet-purified using D Columns on a SuperMACS™ II Separator (Miltenyi Biotec). Magnet-purified parasites were crosslinked in 1% formaldehyde for 10 minutes at 37°C, quenched with glycine to an end concentration of 0.125M and resuspended in FACS buffer (PBS with 2mM HEPES, 2mM glucose, 0.4mM NaHCO3, 0.01% BSA, 2.5mM EDTA). Male and female gametocytes were sorted according to color (GFP for males and RFP for females) on a FACSAria III with a 70uM nozzle at 4°C. Purified gametocyte pellets were lysed in 150ul sonication buffer and chromatin prepared as above.

Ookinetes were cultured *in vitro* as described (Rodríguez et al., 2002). Briefly, mice were pretreated by intra peritoneal injection of 0.2 ml phenylhydrazine (6 mg/ml in PBS) to stimulate reticulocyte formation two days prior to infection with parasites. Gametocyte-enriched blood was harvested via heart puncture and blood was immediately resuspended in 30ml of ookinete medium (RPMI-HEPES complemented with 100uM xanthurenic acid, 200 uM hypoxanthine and 10% BSA, pH7.4) and incubated at 21°C for 24 hours. After 24 hours, ookinetes were magnet-purified using 1ul monoclonal anti-P28 antibody 13.1 (Winger et al., 1988) coupled to 10ul magnetic beads (Dynabeads™ M-280 Sheep Anti-Mouse IgG). Purified ookinetes were crosslinked in 1% formaldehyde for 10 minutes at 37°C, quenched with glycine to an end concentration of 0.125M. Purified ookinetes pellets were lysed in 150ul sonication buffer and chromatin was prepared as above.

### Chromatin immunoprecipitation

Antibodies used for ChIP were rabbit anti H3K9ac (Diagenode Cat# C15410004, RRID:AB_2713905) and rabbit anti-*Pb*HP1 (15). 1ug antibody was incubated with up to 500ng chromatin in ChIP buffer (5% TritonX-100, 750 mM NaCl, 5 mM EDTA, 2.5 mM EGTA, 100 mM Hepes) at 4°C over night. The next day, 50ul Protein A Dynabeads (Fisher 10001D) were added and further incubated for 1-2 h. After washing with buffers containing 100, 150 and 250 mM NaCl, immunoprecipitated DNA was eluted and purified using PCR minelute purification columns (Qiagen).

For each antibody several ChIP reactions were performed in parallel to obtain sufficient amount of DNA for ChIP-seq: (asexual blood stages (3xH3K9ac and 8x *Pb*HP1); female/male gametocytes (3x H3K9ac and 6x *Pb*HP1); ookinetes (2x H3K9ac and 5x *Pb*HP1)).

For each stage, the following numbers of biological replicates were pooled to obtain enough material: ABS (1); FG (5), MG(5), OOK (2).

### ChIP sequencing

For each sequencing library up to 10 ng of ChIP or input DNA were end-repaired, extended with 30 A-overhangs and ligated to barcoded NextFlex adapters (Bio Scientific) as described previously (28). Libraries were amplified (98°C for 2 min; four cycles 98°C for 20 sec, 62°C for 3 min; 62°C for 5 min) using KAPA HiFi HotStart ready mix (KAPA Biosystems) and NextFlex primer mix (Bio Scientific) as described (29). 225-325 bp fragments (including the 125 bp NextFlex adapter) were size-selected using a 2% E-Gel Size Select agarose gel (Thermo Fisher Scientific) and amplified by PCR for ten (asexual blood stages and ookinetes) or eleven (female and male gametocytes) cycles under the same condition as described above. Library purification and removal of adapter dimers was performed with Agencourt AMPure XP beads in a 1:1 library:beads ratio (Beckman Coulter). ChIP-seq libraries were sequenced for 75 bp single-end reads using the NextSeq 500/550 High Output v2 kit (Illumina) on the Illumina NextSeq 500 system.

Sequencing reads were mapped against the *P. berghei* ANKA reference genome v3 using BWA samse (v0.7.12-r1039) (30) (and filtered to mapping quality ≥15 (SAMtools v1.2) (31). Only uniquely mapped reads were used for further analysis. The PCA plot was calculated using default settings in the DeepTools2 suite (32) using non-overlapping 1000bp bins.

### *Pb*HP1 analysis

The bam files containing mapped reads from each ChIP was normalised against its input using *bamcompare* in DeepTools2 (32). For *Pb*HP1, the default settings were used with the following changes: 100bp bin size, 0.01 pseudocount and 1000bp smoothing.

The mean of the log2 *Pb*HP1/input ratio was then extracted for each gene, making each gene fit 500bp using one 500bp bin using *computematrix* in DeepTools2 (32) with default settings with the following changes: missing values = 0 and skip 0. The resulting *bigwig* files were hierarchically clustered with average linkage and Euclidean distance as similarity metric in Cluster3.0 (33). The resulting heatmap and tree was inspected in Treeview (34).

The multigene family list is based on Table S4 from Otto et al, (35), and we updated gene IDs to the *P. berghei* genome version 3 and found 379 genes.

For visualising our data next to (15) we used their *Pb*HP1 over input bedgraph file from the GEO database (GSE102695) and viewed it in IGV (36). Chromosome plots were drawn using Phenogram (http://visualization.ritchielab.psu.edu/phenograms).

### H3K9ac analysis

The bam files containing aligned reads from each ChIP was normalised against its input using *bamcompare* from the *DeepTools2 suite* (32). For H3K9ac, default settings were using with the following changes: bin size 50bp, 0.01 pseudocount, 100bp smoothing. Analysis using heatmaps and summary plots were performed using the deeptools2 (32) suite on usegalaxy.org as well as usegalaxy.eu.

For the ribosomal subunit genes, we used the *P. berghei* orthologues from *P. falciparum* reported in (37). For statistical analysis and drawing of boxplots, we used Graphpad Prism 7.03. Wilcoxon matched-pairs signed rank test was performed on each data set and P-values <0.01 are reported, with (****) P ≤ 0.0001.

### Motif identification

5’ UTRs were extracted from *Pb*ANKA genome version 3. Sequences between annotated genes were attributed to the nearest gene as follows. For head-to-head genes, the 5’ UTRs were split 1:1. For tail-to-head genes, the sequence was split 1:2 (1/3 was assigned as 3’ downstream region to the gene ending and 2/3 was assigned as 5’ UTR to the gene starting). To find motifs enriched in ookinete-expressed genes, we used DREME (v 5.0.0) (38) with default settings locally on the MEME suite (39).

We used the build-in Gene Ontology tool of PlasmoDB with default settings to identify enriched GO terms (40).

### RNA preparation

Asexual blood stage parasites were harvested via heart puncture and resuspended in 15ml RPMI-HEPES and passed through a Plasmodipur filter (Europroxima) to remove white blood cells. Parasite/RBC pellet was lysed with 10ml RBC-lysis buffer (150mM NH_4_Cl, 10mM KHCO_3_, 1mM EDTA) for 20min on ice. Parasite-pellets were washed once in ice-cold PBS and lysed in 500ul TRIzol (Invitrogen) and stored at -80°C.

The method for the purification of gametocytes was modified from (27). Briefly, mice were pretreated by intra peritoneal injection of 0.2 ml phenylhydrazine (6 mg/ml in PBS) to stimulate reticulocyte formation two days prior to infection with parasites. Gametocytes were harvested via heart puncture and blood was immediate resuspended in 4°C coelenterazine loading buffer (CLB) (1x PBS, 20 mM HEPES, 20 mM glucose, 4 mM sodium bicarbonate, 1 mM EDTA, 0.1% BSA in PBS, pH 7.24–7.31) and magnet-purified using D Columns on a SuperMACS™ II Separator (Miltenyi Biotec). Magnet-purified parasites were resuspended in FACS buffer (PBS with 2mM HEPES, 2mM glucose, 0.4mM NaHCO3, 0.01% BSA, 2.5mM EDTA). Male and female gametocytes were sorted according to color (GFP for males and RFP for females) on a FACSAria III with a 100uM nozzle at 4°C. Purified gametocyte pellets were lysed in 500ul TRIzol (Invitrogen) and stored at -80°C.

Ookinetes were cultured *in vitro* as described (Rodríguez et al., 2002). Briefly, mice were pretreated by intra peritoneal injection of 0.2 ml phenylhydrazine (6 mg/ml in PBS) to stimulate reticulocyte formation two days prior to infection with parasites. Gametocytes-enriched blood was harvested via heart puncture and blood was immediately resuspended in 30ml of ookinete medium (RPMI-HEPES complemented with 100uM xanthurenic acid, 200 uM hypoxanthine and 10% BSA, pH7.4) and incubated at 21°C for 24 hours. After 24 hours, ookinetes were magnet**-**purified using 1ul monoclonal anti-P28 antibody 13.1 (Winger et al., 1988) coupled to 10ul magnetic beads (Dynabeads™ M-280 Sheep Anti-Mouse IgG) and resuspened in 500ul TRIzol (Invitrogen) and stored at -80°C.

Genomic DNA was used as a control for the library preparation protocol. For this, asexual blood stage parasites were harvested via heart puncture and resuspended in 15ml RPMI-HEPES and passed through a Plasmodipur filter (Europroxima) to remove white blood cells. Parasite/RBC pellet was lysed with 10ml RBC-lysis buffer (150mM NH_4_Cl, 10mM KHCO_3_, 1mM EDTA) for 20min on ice. Parasite-pellets were washed once in ice-cold PBS and lysed in buffer A (500mM NaAc, 100mM NaCl, 1mM EDTA, pH5.2) and 3% SDS. DNA was extracted using phenol:chloroform and sheared to 150bp fragments on a Bioruptor® Plus sonication device (Diagenode).

### RNA sequencing

RNA was extracted using the Direct-zol™ RNA MiniPrep kit (Zymo). Residual gDNA was digested with the TurboDNA-free kit (Ambion). Stranded RNA sequencing libraries were prepared using the RNA HyperPrep Kit (KAPA) following manufacturer’s protocol with the exception of the amplification step set to 60°C. The RNA library was sequenced in 43bp paired-end reads on a Illumina NextSeq 500 sequencer and each sample was split over four non-independent lanes.

RNAseq was performed using biological triplicates for each condition.

We used sheared genomic DNA (gDNA) as a technical control and did not observe a bias towards GC-rich(er) sequences (**Figure S2A**), a known problem in *Plasmodium falciparum* NGS (14).

### RNA sequencing analysis

Quality of the reads were checked by eye using FASTQC. Reads were aligned separately for each lane to *Pb*ANKA genome v3 using HiSat2 with default settings (v 2.0.5.2; --max intron size 5000; -- fr) (41). Mapped reads were pooled into one bam file per condition and replicate. Importantly, to adjust for potential GC-bias gDNA was processed alongside RNA samples as a control for the RNA library preparation. GC-bias was calculated according to (42) using the default settings for *computeGCBias* in DeepTools2 suite (32).

The PCA plot was calculated using default settings in the DeepTools2 suite (32).

We calculated the FPKM (Fragments Per Kilobase of transcript per Million mapped reads) value for each gene using featurecounts with default settings (43).

For comparative transcriptomics we used Deseq2 (v2.11.39) with default settings (44). We considered significantly differently regulated genes as having a p value of lower than 0.001 (adjusted for multiple testing with the Benjamini-Hochberg procedure which controls false discovery rate (FDR)).

All of the analysis has been performed using usegalaxy.org (an open source, web-based platform for data intensive biomedical research) (45) unless stated otherwise. Additionally, usegalaxy.eu was used for some calculations.

## RESULTS

### Generation of epigenetic and transcriptional profiles

ChIP was carried out on *P. berghei* mixed ABS, FG, MG and OOK using antibodies against *P. berghei* HP1 (*Pb*HP1) (15) and H3K9ac. To prevent sample contamination, ABS were sampled from the non-gametocyte producer ANKA 2.33 parasite line (24). MG and FG were isolated from the *820cl1m1cl1* (wt-fluo) line (26) using flow cytometry sorting. Purified OOK were prepared from *in vitro* cultures of the *507m6cl1* line (25), 24 hours post gametocyte activation. Immuno-precipitated DNA was amplified and subjected to next-generation sequencing (NGS). Principal component analysis (PCA) showed that H3K9ac and *Pb*HP1 samples are clustered together and separately from each other, respectively, suggesting that euchromatin and heterochromatin occupancies differ from each other but are similar between the three stages and parasite lines (**Figure 1A**).

**Figure 1.**
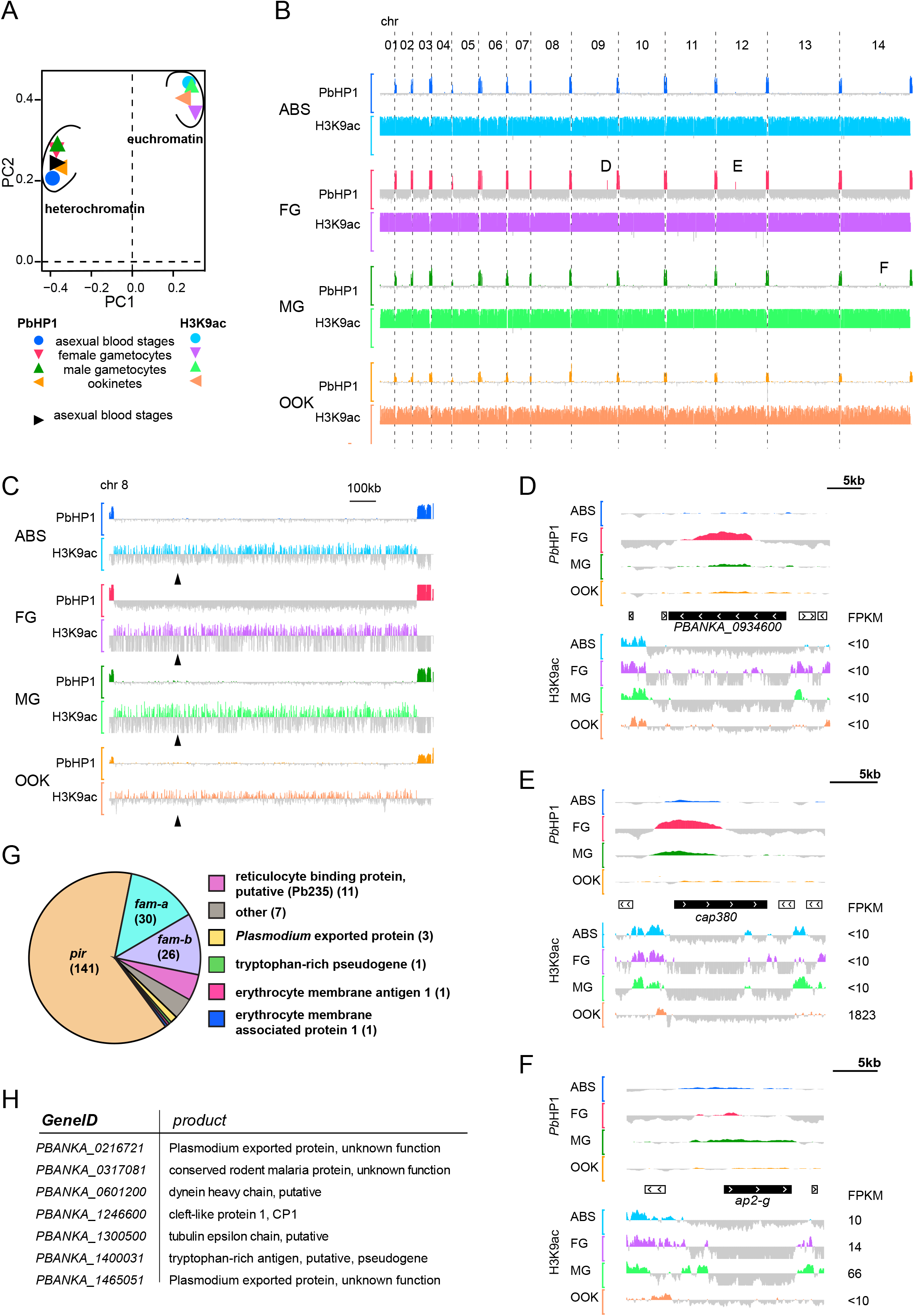
Heterochromatin distribution remains stable during malaria parasite development whereas euchromatin distribution is dynamic. (A) Principal component analysis of log_2_ transformed ChIP over input data. Heterochromatin and euchromatin cluster away from each other. Data from ABS from a previous heterochromatin study is included (15). (B) Screenshot of heterochromatin and euchromatin distribution across all 14 *P. berghei* chromosomes for each developmental stage. Peaks correspond to log_2_ transformed data of either *Pb*HP1-ChIP/input or H3K9ac-ChIP/input. Log_2_ scale for PbHP1 is (−3 to 3) and (−2 to 2) for H3K9ac, respectively. Letters highlight chromosome-central genes showing enrichment in *Pb*PH1 binding. Their close-up view is shown in Fig. 1D-F. (C) Bar chart close-up view of heterochromatin and euchromatin occupancy in chromosome 8. The approximate location of the centromere (syntenically inferred from *P. falciparum* (46)) is indicated with arrowheads. (D-E) Chromosome central genes associated with heterochromatin in this study. (F) *ap2-G* does not qualify as heterochromatic in our study. Black boxes show gene of interest, white boxes indicate neighbouring genes and chevrons indicate the transcriptional orientation for each gene. Peaks correspond to log_2_-transformed data of *Pb*HP1-ChIP/input (−3 to 3) and H3K9ac-ChIP/input (−2 to 2). (G) Pie chart of identified heterochromatic genes (including pseudogenes) grouped into gene families as described previously (35). The number of genes in each family is shown in parentheses. (H) Description of seven heterochromatic genes not belonging to known multigene families. ABS, asexual blood stages; FG, female gametocytes; MG, male gametocytes; OOK, ookinetes

To investigate the link between epigenetic profiles and gene expression, we complemented the epigenetic profiles with transcriptomics data from the same stages and parasite lines by RNAseq. PCA analysis showed that transcription profiles of ABS and MG are more similar to each other than to the other two stages, and that transcription profiles of FG are more related to OOK (**Supplementary Figure 1**).

### Mutually exclusive profiles of heterochromatin and euchromatin

Bar plots of ChIPped vs input chromatin showed that heterochromatin is confined to telomeric and subtelomeric regions of all chromosomes, except from the right arm of chromosome 4, in contrast to H3K9ac that is detected in all other chromosomal regions (**Figure 1B**). Ten subtelomeric regions could only be partially mapped due to their repetitive nature. As shown in the example of chromosome 8 (**Figure 1C**), but observed for all chromosomes and in all stages, centromeres do not show enrichment in heterochromatic marks (46). Interestingly, *P. berghei* chromosomes are largely devoid of chromosome internal heterochromatin islands, in contrast to most other *Plasmodium* species (15).

To identify heterochromatic genes we calculated the mean *Pb*HP1 occupancy (log_2_ ratio of *Pb*HP1 ChIP to input) of the gene open reading frame (ORF) and performed hierarchical clustering. Only two non-telomeric and non-subtelomeric genomic regions showed clear heterochromatic profiles. The first region encompasses *PBANKA_0934600*, a gene encoding a large, conserved protein of unknown function with orthologues in all *Plasmodium* species (**Figure 1D**). This region is heterochromatic in all four stages investigated. Transcript levels of *PBANKA_0934600* are very low (Fragments Per Kilobase of transcript per Million mapped reads (FPKM) <10) suggesting that the gene is not expressed in any of the stages investigated here (**Supplementary Table 1**). Proteomic data indicate that its *P. falciparum* ortholog (*PF3D7_1113000*) is expressed in sporozoites (47). Nevertheless, a previous study showed that *PBANKA_0934600* is redundant for *P. berghei* transmission (48).

The second chromosome-central heterochromatic region corresponds to *cap380* and appears to be epigenetically silenced in ABS, FG and MG but not in OOK (**Figure 1E**). Indeed, OOK display increased levels of relative *cap380* transcripts, which coincide with a sharp H3K9ac peak within the *cap380* 5’UTR. *Cap380* transcription is controlled by AP2-O (23, 35) and the protein is expressed in early-stage oocysts and localizes to the oocyst capsule (49, 50). Our data provide evidence that heterochromatic silencing is an additional regulatory level of *cap380* expression, presumably preventing its premature transcription.

A third region that showed clear heterochromatic marks by visual inspection but could not be classified as heterochromatic by our analysis algorithm encompasses the gene encoding AP2-G (*PbANKA_1437500*; **Figure 1F**). Expression of *P. falciparum* AP2-G is epigenetically controlled by *Pf*HP1 (51). The weak levels of *Pb*HP1 marking of this region in our analysis could be explained by our choice of the ANKA 2.33 parasite line. This line carries a mutation in the *ap2-g* gene resulting in expression of a truncated, non-functional protein unable to induce gametocytogenesis (21), thereby making its epigenetic silencing redundant.

### Epigenetic silencing of subtelomeric multigene families

Our analysis revealed that as many as 221 genes (including pseudogenes) located in subtelomeric regions are significantly enriched in *Pb*HP1 binding (**Figure 1G, Supplementary Table 1**). Of these, 214 belong to the rodent malaria parasite (RMP) multigene family (35). It has been previously established that *Plasmodium* heterochromatin is largely associated with gene families involved in antigenic variation and host-parasite interactions (7, 15). Out of the 214 heterochromatic RMP genes, 141 belong to the *pir* (*Plasmodium* interspersed repeat) family that comprises 200 genes and pseudogenes. It is the largest multigene family in *P. berghei* and its members show variegated expression across stages and between acute and chronic infections (35, 52, 53). In *P. chabaudi*, changes in expression of the orthologues *cir* gene family have been associated with parasite virulence (10).

Thirty and 26 of the identified heterochromatic genes belong to the *fam-a* and *fam-b* families, respectively. Members of the *fam-a* family have a steroidogenic acute regulatory-related lipid transfer (START) domain and can transfer phosphatidylcholine *in vitro*, suggesting that these proteins are involved in lipid transport for membrane synthesis (53). Most *fam-b* members have a PYST-B domain, a signal peptide and a PEXEL motif as well as a transmembrane domain (35). Proteins encoded by the *pir, fam-a* and *fam-b* families are exported into the iRBC (54). While the *fam-a* family is present in all *Plasmodium* species, the *fam-b* family is specific to rodent malaria parasites.

Eleven of the heterochromatic, subtelomeric genes belong to the putative reticulocyte binding protein family (aka *Pb235*) (55), and three of them encode unknown *Plasmodium* exported proteins. Finally, erythrocyte membrane antigen 1, erythrocyte membrane associated protein 1 and a tryptophan-rich protein pseudogene are also heterochromatic.

Only seven subtelomeric heterochromatic genes do not belong to RMP multigene families (**Figure 1H**). They include: *PBANKA_1246600* that encodes CP1, an atypical PEXEL protein exported to discrete structures in the cytosol of iRBCs (56); *PBANKA_0216721* and *PBANKA_1465051* both encoding exported proteins of unknown function; *PBANKA_0317081* encoding a conserved rodent malaria parasite protein; *PBANKA_1300500* encoding a putative tubulin epsilon chain; *PBANKA_1400031*, a putative pseudogene encoding a tryptophan-rich antigen; and *PBANKA_0601200* encoding a dynein heavy chain.

*PBANKA_0601200* is one of 22 dynein-related proteins annotated in the *P. berghei* genome (**Figure S2**). Dyneins are one of three cytoskeletal motor protein families in eukaryotes and made up of a protein complex of heavy, light and intermediate chain. Intriguingly, *PBANKA_0601200* is the only heterochromatic dynein gene in this study but not orthologous to neither of two heterochromatic dynein heavy chain-encoding genes found in *P. falciparum* blood stages (Flueck et al., 2009). Our data show that the gene is transcribed in MG, and its promoter is rich in H3K9ac binding in gametocytes (**Figure S2**). Its knockout is associated with slow ABS growth (57). It is interesting that of six dynein genes located close to telomeres, only *PBANKA_0601200* is found to be heterochromatic in our study.

In summary, our data show that similar to other *Plasmodium* species (15), subtelomeric multigene families are epigenetically silenced via *Pb*HP1 throughout *P. berghei* development. Interestingly, and in contrast to *P. falciparum* (15), we did not observe any expansion of heterochromatic boundaries throughout the *P. berghei* lifecycle.

### Variant developmental expression of heterochromatic genes

Stochastic changes in heterochromatin distribution result in clonal variant gene expression, which forms the basis of *P. falciparum* antigenic variation (6). Since different parasite lines were used in our study (except FG and MG that derived from the same line), we could not directly determine whether differences in heterochromatin distribution are due to the line epigenetic background or developmental stage, nonetheless, comparative heterochromatic profiling would capture the sum of both. We compared the heterochromatic profiles of genes across development, and found a surprisingly small number of 16 genes with differential heterochromatin occupancy between stages (**Figure 2A** and **Supplementary Figure S3**). This small number stands in sharp contrast to the 252 genes shown to exhibit clonal variant expression in *P. falciparum* ABS alone (58). Thirteen of the 16 genes belonged to multigene families, and 12 showed *Pb*HP1 enrichment in FG but not in MG. Since both gametocyte samples were derived from the same line (26), these data could indicate true developmental epigenetic differences that in turn could imply that heterochromatin occupancy is reorganized during gametocyte development. However, transcript abundance of these 12 genes was very low (<10 FPKM) in all lifecycle stages examined (**Supplementary Table 1**), suggesting that absence of *Pb*HP1 occupancy may not necessarily result in an active transcriptional state.

**Figure 2.**
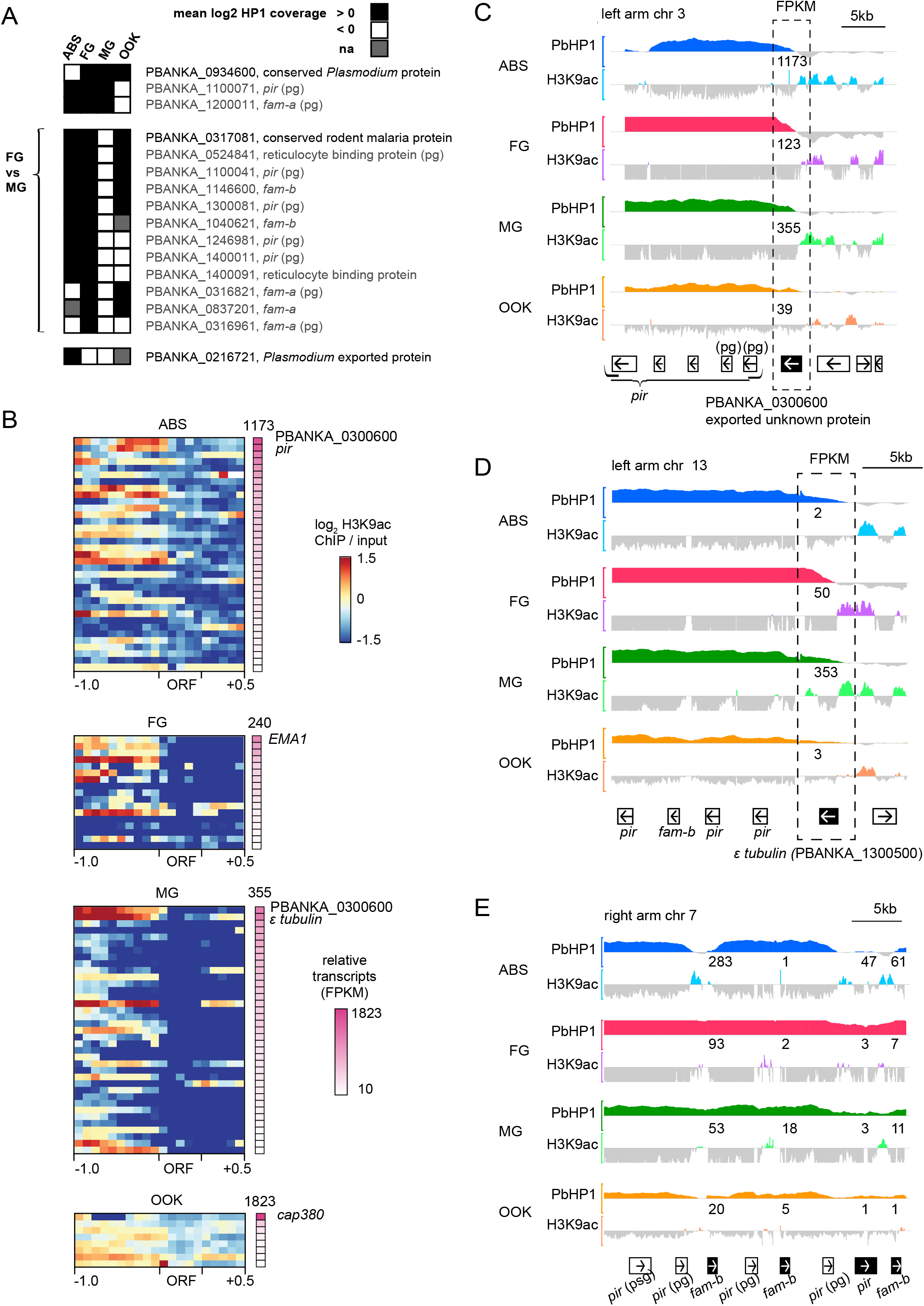
A subset of subtelomerically located genes shows signs of variant expression. (A) Genes with varying *Pb*HP1-occupancy through parasite development. Genes not belonging to a multigene family are highlighted with a darker colour. (B) Heatmap of euchromatic traits (H3K9ac-ChIP/input) of heterochromatic genes with at least 10 FPKM (fragments per kilobase of transcript per million mapped reads) for the developmental stage shown. H3K9ac enrichment for each gene locus is shown as log_2_ transformed H3K9ac ChIP/input (1000bp upstream of ATG the ORF, and 500bp downstream of stop codon, respectively). Pink colour indicates mean relative transcripts (in FPKM), and the highest transcript(s) for each life cycle stage is/are named. Higher transcript numbers do not always correlate with H3K9ac occupancy within 1000bp upstream of the start codon. A list of all genes is found in Table S1. (C) *PbANKA_0300600*, an exported protein of unknown function located at the heterochromatic boundary shows both heterochromatic and euchromatic traits and is transcribed in all four life cycle stages. (D) *PbANKA_1300500*, the tubulin epsilon chain is both heterochromatic and euchromatic in gametocytes, and is most transcribed in male gametocytes. (E) Four genes within the heterochromatic region at the right arm of chromosome 7 display euchromatic traits correlating with transcription. Relative transcripts (in FPKM) for each gene are shown in numbers above ChIP-seq tracks. Black boxes show gene of interest, white boxes indicate neighbouring genes and arrows indicate the transcriptional orientation for each gene. Peaks correspond to log_2_-transformed data of *Pb*HP1-ChIP/input (−3 to 3) and H3K9ac-ChIP/input (−2 to 2). ABS, asexual blood stages; FG, female gametocytes; MG, male gametocytes; OOK, ookinetes; pg, pseudogenes.

To identify if variegated clonal expression is present in *P. berghei* stages, we arbitrarily selected all heterochromatic genes with more than 10 FPKM (mean of three replicates) in at least one developmental stage. Using this approach, we identified an additional 35 genes in ABS, 17 genes in FG, 37 genes in MG and 8 genes in OOK, respectively, which are likely variantly expressed (**Figure 2B**). Most of these heterochromatic genes display additional euchromatic H3K9ac marks, mainly in their 5’UTR and some of them show very high transcript levels (**Figure 2B**). For example, despite the *PbANKA_0300600* ORF being heterochromatic, its 5’UTR is enriched in H3K9ac and the gene is highly expressed in all stages (**Figure 2C**). This suggests that *PbANKA_0300600* is expressed in only a subset of cells. Similarly, the epsilon-tubulin ORF is heterochromatic in all stages but shows euchromatic traits and high transcript levels in both MG and FG (**Figure 2D**). The function of epsilon-tubulin in *Plasmodium* is unknown, but the protein is known to mark the older of the two human centrioles upon centrosome duplication (59) and be an essential part of the basal bodies in *Tetrahymena* (60).

Another example of clonally variegated expression includes multigene family members on the right arm of chromosome 7, displaying both heterochromatin and euchromatin occupancies and high transcript levels (**Figure 2E**). This is consistent with the finding that some heterochromatic genes are transcribed by a subset of cells, effectively displaying variegated expression as shown before for members of putative exported protein families in *P. berghei* (Fougère et al., 2016; Pasini et al., 2013). Genes displaying hallmarks of clonally variant expression are all located at the heterochromatin-euchromatin boundaries, which seems to facilitate clonal variation.

### Distinct H3K9ac distribution in ribosomal protein genes

H3K9ac is a universal histone mark associated with active promoters across the animal kingdom, including *P. falciparum* ABS, oocysts and sporozoites (12, 61, 62). We investigated the relationship between H3K9ac distribution and gene transcription across the three *P. berghei* developmental stages. H3K9ac enrichment in the gene ORF, 1 kb 5’UTR and 500 bp 3’UTR was examined against transcript abundance for each stage (**Figure 3A**). In consistence with previous findings from *P. falciparum*, a positive correlation was detected between the gene 5’UTR H3K9ac enrichment and transcriptional levels in ABS (12, 13). We also found that H3K9ac enrichment in the 5’UTR positively correlates with transcription levels in MG and OOK but less so in FG (**Figure 3B**).

**Figure 3.**
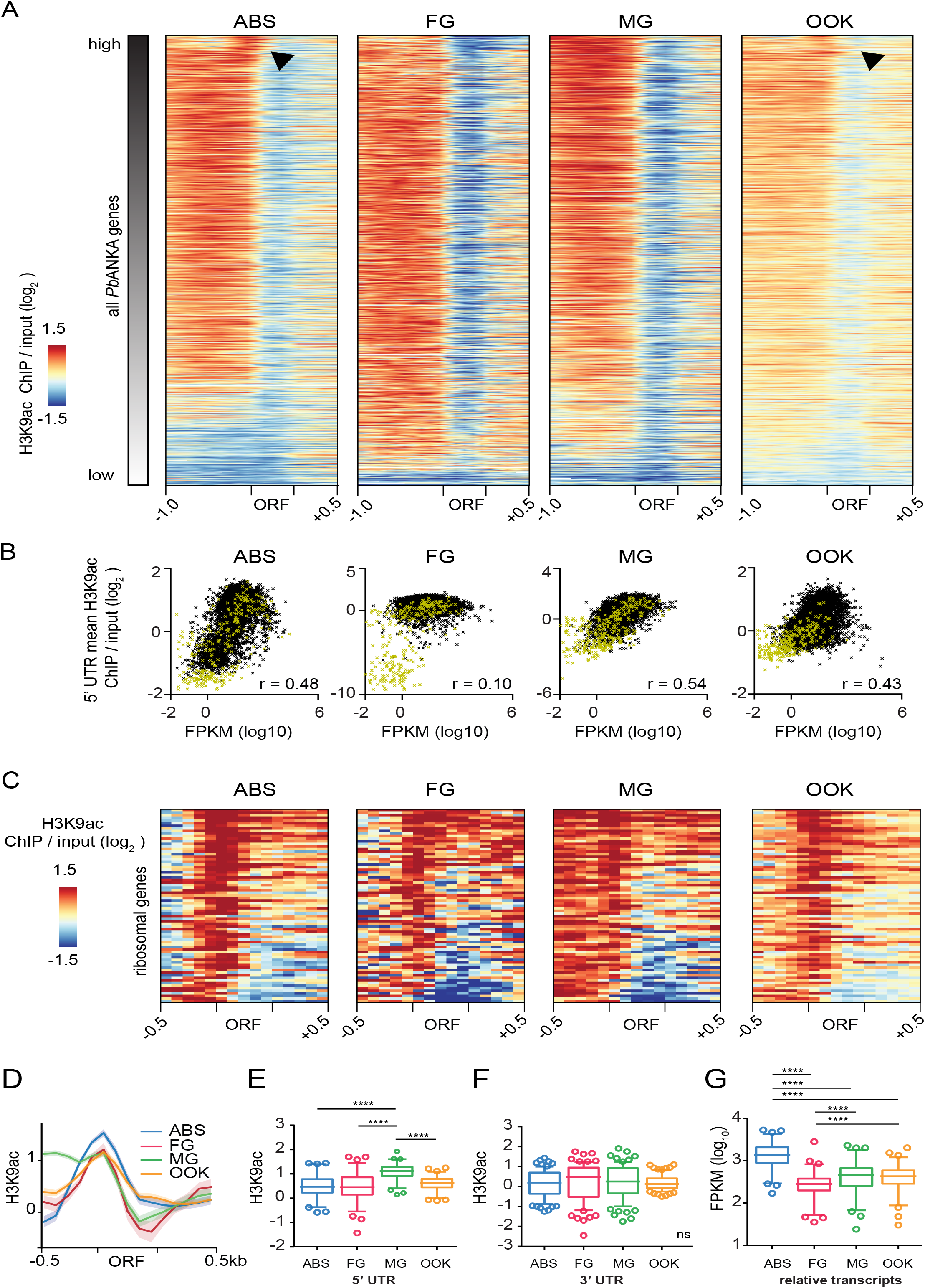
5’ UTRs of ribosomal protein genes display a distinctive H3K9ac pattern. (A) Heat map of H3K9ac distribution in malaria parasite development. Genes at each stage are sorted according to their relative transcription levels, with highly expressed genes on top. Arrowheads indicate a shift of H3K9ac occupancy towards the start codon. H3K9ac enrichment for each gene locus is shown as log_2_-transformed H3K9ac ChIP/input (1000bp upstream of ATG the ORF, and 500bp downstream of stop codon, respectively). (B) Scatterplot showing mean H3K9ac occupancy of 5’ UTRs (1000bp) against transcriptional strength (mean FPKM) for each gene for each developmental stage. Spearman’s rank correlation coefficient for each scatterplot is shown (r). Genes belonging to a multigene family are highlighted in yellow. (C) Heat map of H3K9ac enrichment for all 74 ribosomal protein genes (40S and 60S). H3K9ac is enriched in a sharp peak around the start codon in ABS, FG and OOK, but less in MG. (D) Summary plot of the data shown in C. The mean of H3K9ac enrichment of ribosomal protein genes is shown as a bold line for each life cycle stage, standard error is shown in lighter colours. (E) Boxplots show mean H3K9ac-enrichment values for 5’UTRs (500bp) of ribosomal protein genes for each stage. Whiskers indicate the 5^th^ and 95^th^ percentile, respectively. Individual symbols represent outliers. Asterisks mark significance (Wilcoxon signed rank test for matched-pairs, p≤0.0001). (F) Boxplots show mean H3K9ac-enrichment values for 3’UTR of each ribosomal protein gene. Whiskers indicate the 5^th^ and 95^th^ percentile, respectively. Individual symbols represent outliers. Wilcoxon signed rank test for matched-pairs found no difference between the developmental stages (ns). (G) Boxplots show mean relative ribosomal gene transcripts for each life cycle stage. Asterisks mark significance (Wilcoxon signed rank test for matched-pairs, p≤0.0001). ABS, asexual blood stage; FG, female gametocyte; MG, male gametocyte; OOK, ookinete; ORF, open reading frame

A shift of H3K9ac occupancy toward the gene ORF was observed for highly expressed genes, mainly in ABS and OOK (see arrowheads in **Figure 3A**). Many of the genes with high expression encode ribosomal proteins. Closer investigation indicated that H3K9ac occupancy sharply peaks around the start codon of ribosomal protein genes in ABS, FG and OOK (**Figure 3C**). Intriguingly, H3K9ac occupancy was extended further into the 5’UTR of MG resulting in no clear peak being detected around the start codon (**Figure 3D**). Further analysis confirmed that the mean H3K9ac enrichment in the 500 bp 5’UTR of ribosomal genes was significantly different between MG and all other stages (Wilcoxon signed rank test, p≤0.0001), but no difference was detected between any of the other stages (**Figure 3E**). As a control, the same analysis using the 500 bp 3’UTR of these genes did not show any significant differences between any of the stages (Wilcoxon signed rank test, p>0.3) (**Figure 3F**). Interestingly, relative transcripts of ribosomal protein genes differ between all developmental stages, with highest transcript numbers found in ABS (**Figure 3G**). We did not detect a correlation between transcriptional abundance and H3K9ac occupancy. These data clearly show that genes encoding ribosomal proteins exhibit a different H3K9ac pattern from other genes, highlighting interesting differences in epigenetic makeup for a specific gene group. It has been previously suggested that a shift in peak shape can indicate a different function of a gene (63), and highly dense and narrow distributions of H3K9ac near transcriptional start sites have been associated with constitutive expression of genes involved in translation in plants (64).

### H3K9ac enrichment does not correlate with transcript levels in FG

We further wanted to examine the relationship between H3K9ac occupancy and transcript levels in *P. berghei* development. For this, we identified genes that are differentially expressed between two related developmental stages and examined their H3K9ac occupancy for each of the two stages (**Figure 4A, B, C**). Genes upregulated in ABS (1426), MG (1523) or OOK (1641) compared to FG exhibit greater H3K9ac enrichment in their 5’UTR (1000bp upstream of the start codon) in each respective stage than genes downregulated in any of these three stages (1429, 1518 and 1534, respectively) compared to FG (**Figure 4D, E, F**). However, when the same groups of genes were examined for H3K9ac enrichment in their 5’UTR in FG, upregulated genes in FG showed less (FG vs. ABS and FG vs. MG) or the same (FG vs. OOK) H3K9ac enrichment than downregulated genes (**Figure 4G, H, I**). These results indicate that H3K9ac occupancy in the gene 5’UTR is a good predictor for relative transcript levels in ABS, MG and OOK.

**Figure 4.**
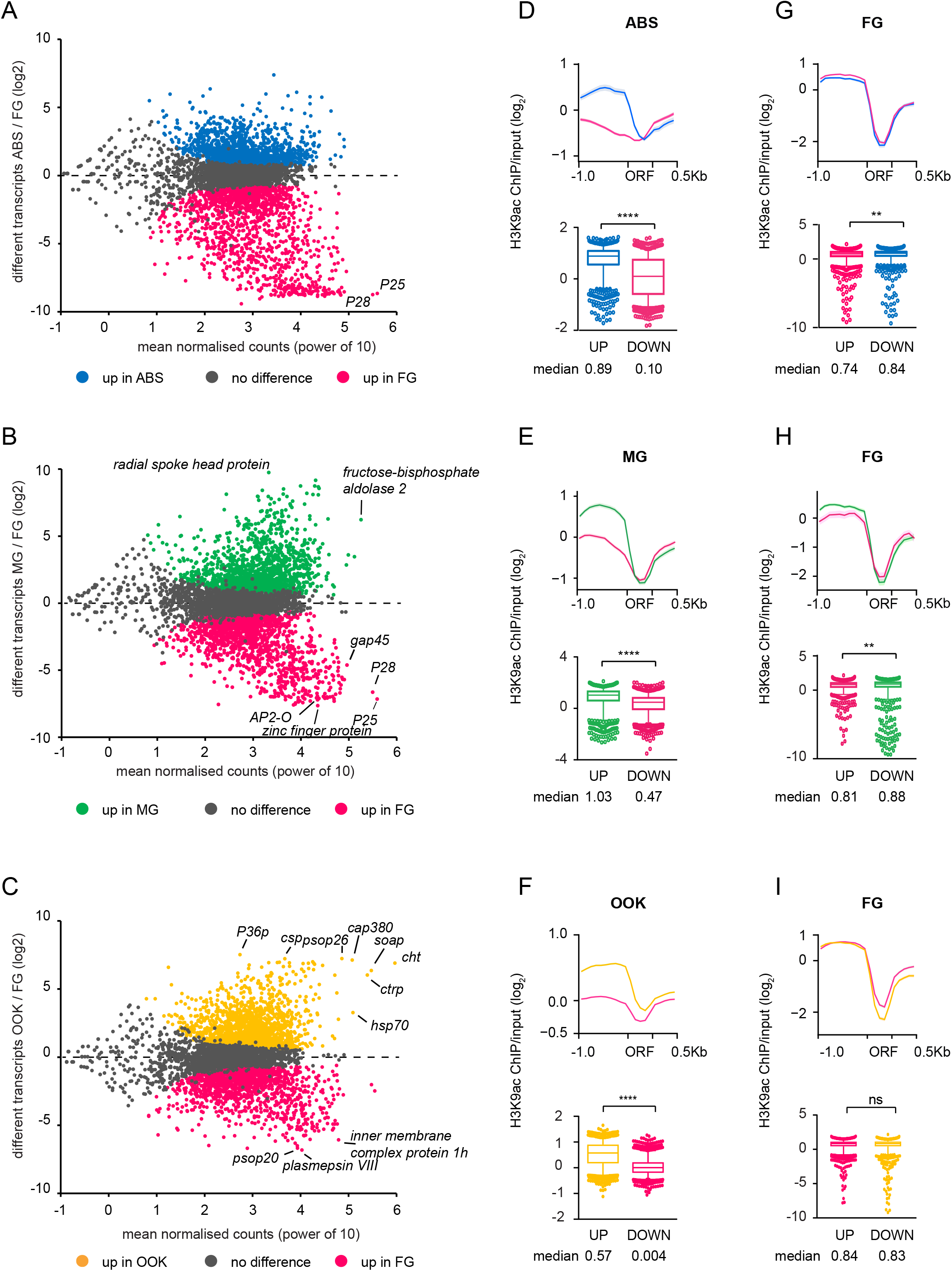
H3K9ac intensity correlates with relative transcripts in asexual blood stages, male gametocytes and ookinetes. (A-C) Comparative profiling of ABS, MG and OOK vs FG. Each dot represents a gene. Genes with a non-significant change in transcript abundance are shown in grey and genes that are significantly up- or downregulated are shown in their respective colour (p value adjusted for multiple testing with the Benjamini-Hochberg procedure which controls false discovery rate (FDR), here p<0.001). (D) Mean H3K9ac distribution of in asexual blood stages correlates with transcript levels. The upper panel shows the mean H3K9ac distribution for all genes that are either up- or downregulated in ABS vs FG. The mean H3K9ac enrichment of all loci is indicated with a solid line, and the standard error is shown in a lighter colour. 1000bp upstream and 500bp downstream of each gene is included. Boxplots show mean H3K9ac enrichment for the 5’ UTR for both gene groups. In ABS, (compared to FG), mean H3K9ac enrichment in the 5’UTR of a gene positively correlates with its transcript levels (Mann Whitney test, p<0.0001). Boxes extend from the 25th to 75th percentiles, and the median is shown as a line in the middle of the box and as a number below the boxes. Whiskers indicate the 5^th^ and 95^th^ percentile, respectively. Individual symbols represent outliers. The median is shown. (E) In male gametocytes (compared to FG), mean H3K9ac enrichment in the 5’UTR of a gene positively correlates with its transcript levels (Mann Whitney test, p<0.0001). Labelling is the same as in D. (F) In ookinetes (compared to FG), mean H3K9ac enrichment in the 5’UTR of a gene positively correlates with its transcript levels (Mann Whitney test, p<0.0001). Labelling is the same as in D. (G) Mean H3K9ac distribution of in female gametocytes using the same gene groups as in A and D. Genes that are upregulated in FG (compared to ABS) are less enriched for H3K9ac in their 5’UTR of than downregulated genes in FG (Mann Whitney test, p=0.0016). (H) Mean H3K9ac distribution of in female gametocytes using the same gene groups as in B and E. Genes that are upregulated in FG (compared to MG) are less enriched for H3K9ac in their 5’UTR of than downregulated genes in FG (Mann Whitney test, p=0.0024). (J) Mean H3K9ac distribution of in female gametocytes using the same gene groups as in C and F. Genes that are upregulated and downregulated in FG (compared to OOK) show the same H3K9ac enrichment in FG (Mann Whitney test, non-significant (ns), p= 0.57).

We further investigated the absence of positive correlation between H3K9ac and transcript levels by generating two high-confidence lists of genes that are either upregulated (591 genes) or downregulated (271 genes) in FG compared to all other stages, respectively (**Figure 5A** and **Supplementary Table 3**). Gene Ontology (GO) analysis using the GO slim version showed that the former list includes genes involved in cell adhesion, reproduction, motility, differentiation, cell cycle and locomotion, all of which are characteristic of cells preparing for fertilization and development into the motile ookinete stage (**Figure 5B** and **Supplementary Table 2**). In contrast, downregulated genes in FG are involved in ribosome biogenesis and translation, suggestive of a translationally less active FG stage compared to the other three stages. FG are known to produce and store mRNAs that are translationally repressed (2). Similar to before, these downregulated FG genes show significantly higher H3K9ac enrichment in their 5’UTRs than genes that are upregulated in FG (**Figure 5C**).

**Figure 5.**
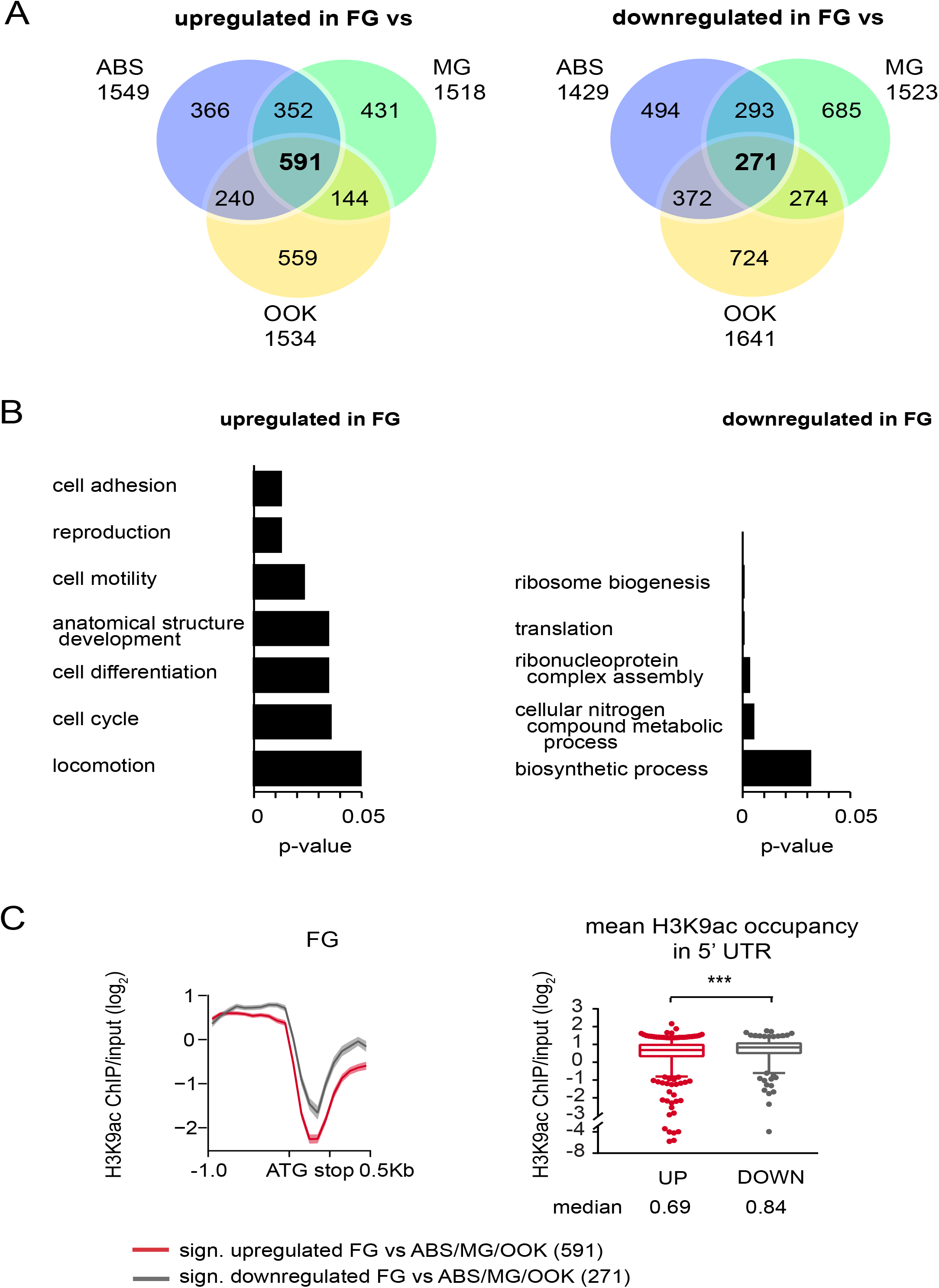
H3K9ac enrichment in the 5’UTR of a gene does not positively correlate with its relative transcripts in female gametocytes. (A) Venn diagram of all genes that are significantly up- or downregulated in female gametocytes compared to asexual blood stages, male gametocytes and ookinetes, respectively. (B) Enriched GO-terms (slim) of both up- and downregulated genes in female gametocytes compared to all other life cycle stages. Each respective p-value is indicated. The full list of genes and GO-terms can be found in Supplementary Table 3. (C) H3K9ac enrichment in female gametocytes for female-specific genes. The left panel shows H3K9ac enrichment in female gametocytes for genes that are significantly up- or downregulated in female gametocytes only. The mean H3K9ac enrichment of all loci is indicated with a solid line, and the standard error is shown in a lighter colour. 1000bp upstream and 500bp downstream of each gene is included. Boxplots show mean H3K9ac enrichment for the 5’ UTR for both gene groups. Mean H3K9ac enrichment in the 5’UTR of a gene negatively correlates with its transcript levels (Mann-Whitney test, p-value=0.0009). Boxes extend from the 25th to 75th percentiles, and the median is shown as a line in the middle of the box. Whiskers indicate 5 and 95 percentiles, respectively, and outliners are individual dots. The median is shown.

These data together strongly indicate that H3K9ac, a histone mark otherwise universally associated with active promoters, does not directly correlate with transcript levels in FG. Instead, it appears that H3K9ac marks the promoters of most genes in FG irrespective of their transcript levels. Furthermore, genes that are translationally repressed by the DOZI-complex show lower H3K9ac occupancy compared to all euchromatic genes, corroborating that high transcript levels do not positively correlate with H3K9ac in FG (**Supplementary Figure S5A-B**). Also, H3K9ac enrichment in FG is unrelated to future transcriptional activity, as AP2-O regulated genes (expected to be active in OOK) (23) display the same H3K9ac enrichment compared to all euchromatic genes in the *P. berghei* genome (**Supplementary Figure S5C-D**). It is therefore possible that FG is a stage where epigenetic remodeling takes place, and that this remodeling includes universal marking of gene promoters with H3K9ac.

### Novel and known DNA motifs control ookinete gene expression

In most metazoans as well as in flowering plants, maternally deposited proteins and mRNAs are responsible for directing early stages of development post fertilization, while *de novo* transcription is resumed via specific transcription factors, a phase called maternal-to-zygotic transition (MZT). In *Plasmodium*, maturation of the fertilized female gamete to OOK is dependent upon both de-repression of maternal mRNAs (2, 65) and *de novo* transcription mediated by ookinete-specific transcription factors such as AP2-O (16, 17, 23).

We further investigated the mechanisms of MZT in *P. berghei* starting with genes that are differentially upregulated in OOK compared to FG (**Figure 4C**), and compared them to 464 genes previously annotated as controlled by AP2-O (23) (**Figure 6A**). Of the 1641 ookinete-specific genes, only 302 qualified as AP2-O controlled (**Figure 6A**), suggesting that not all AP2-O controlled genes have been annotated to date and/or that additional transcription factor(s) might be involved in ookinete-specific transcription. Therefore, we searched for DNA-motifs (38) that are significantly enriched in the 5’UTRs of all 1641 OOK-enriched genes and found two highly significantly enriched motifs: the AP2-O CTAGCT/CA motif (17) present in 265 genes, and the AAAAAAAA motif found in as many as 1571 genes (**Figure 6A**).

**Figure 6:**
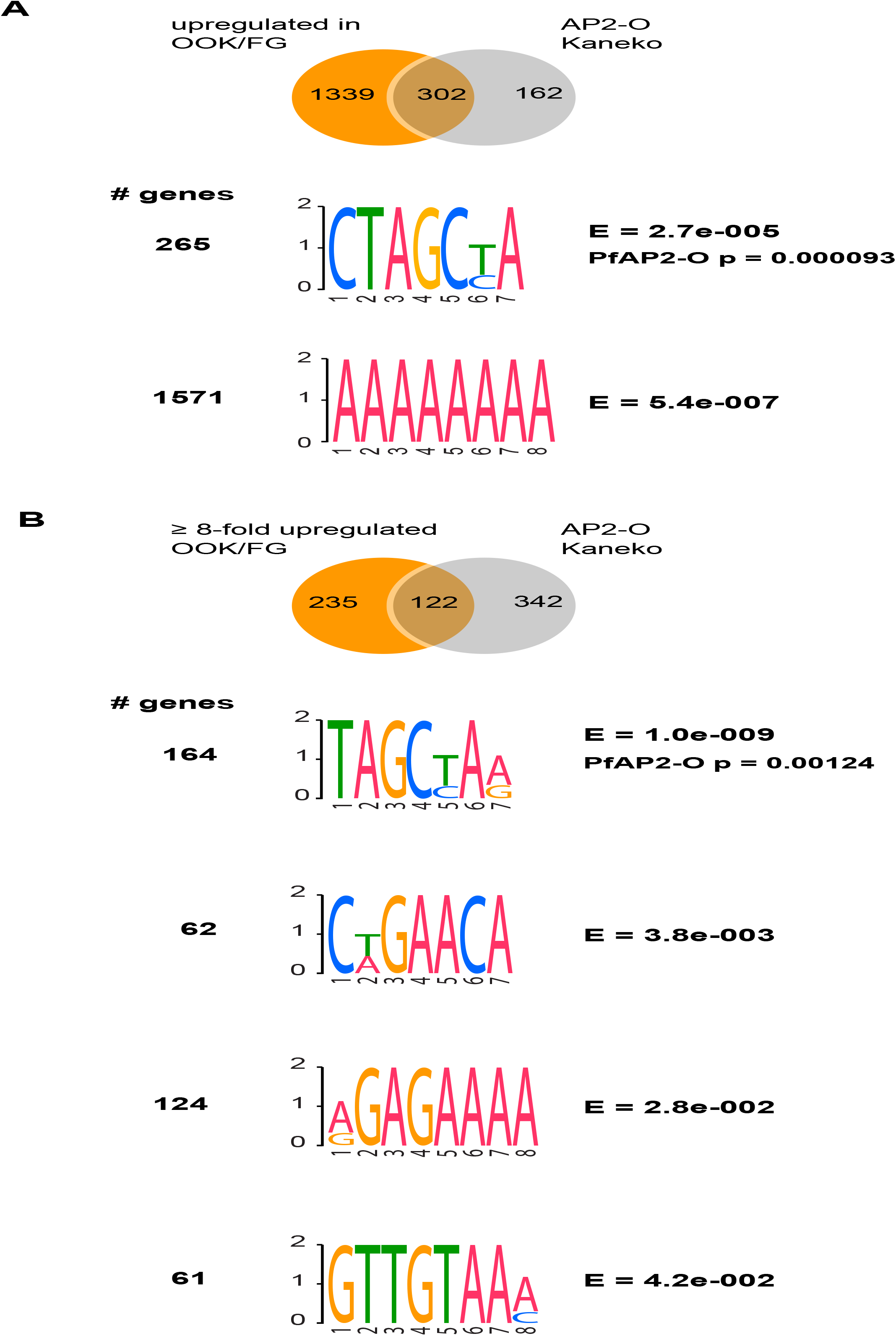
Novel and known DNA-binding motifs enriched in the 5’ UTR of ookinete-specific genes. (A) Venn diagram of genes that are significantly up regulated in ookinetes vs female gametocytes and their overlap with AP2-O genes from Kaneko et al (23).DNA motifs significantly enriched in in the 5’ UTR of all significantly upregulated ookinete genes compared to the 5’ UTR of all genes. The number of genes exhibiting each motif is shown as well as the E-value for each motif. The AP2-O motif is highly enriched. (B) Venn diagram of genes that are at least 8-fold significantly up regulated in ookinetes vs female gametocytes and their overlap with AP2-O genes from Kaneko et al (23) DNA motifs significantly enriched in the 5’ UTR of 8-fold upregulated ookinete genes compared to the 5’ UTR of all genes. The number of genes exhibiting each motif is shown as well as the E-value for each motif. The AP2-O motif is highly enriched.

We repeated the same search on a more stringent list of 357 genes that are at least 8-fold upregulated in OOK compared to FG; and only 122 qualified as controlled by AP2-O (**Figure 6B**). Four DNA-motifs were identified, the most significant of which again resembles the PfAP2-O motif (TAGCT/CAA/G) and was found in almost half of all genes (164). The other three novel motifs were CT/AGAACA (in 62 genes), A/GGAGAAAA (in 124 genes) and GTTGTAAA/C (in 61 genes).

These data confirm previous findings that AP2-O is a master regulator of *de novo* transcription and MZT in *Plasmodium* and indicate the existence of additional transcription factors involved in this process. It is important to note that while additional OOK transcription factors have been described (16), none of the published DNA-motifs matches those found here.

## DISCUSSION

Our study provides new insights into the epigenetic regulation of *Plasmodium* gene expression during its developmental transition from asexual to sexual and zygotic stages, which occurs as the parasite moves from its vertebrate host to the insect vector and is essential for parasite transmission.

Epigenetic silencing has been associated with key strategies of parasite development and environmental adaptations. Our data reveal that HP1-mediated epigenetic silencing remains largely unchanged during this transition and confined to chromosomal subtelomeric regions. Mapping *Plasmodium* subtelomeric genes is very challenging as most genes belong to multigene families with high levels of sequence identity between them. Indeed, as many as 51 heterochromatic genes could not be mapped to *P. berghei* chromosomes in our analysis: 14 *fam-a*, 8 *fam-b*, 1 fam-c, 27 *pir*, and one encoding a tryptophan-rich antigen.

Comparison of our findings with those of Fraschka et al. (15) that analyzed *Pb*HP1 occupancy in ABS of the parental *P. berghei* ANKA strain revealed that 177 heterochromatic genes are shared between the two studies, 46 are specific to our study and 14 are specific to the other study (**Supplementary Figure S4A**). Whereas the vast majority of differences pertain to subtelomeric regions, where our analysis appears to be more sensitive, the combination of the two studies identified four non-subtelomeric heterochromatic genes, of which only *cap380* encoding an oocyst capsule protein is detected in both studies. On the one hand, *PBANKA_0934600* that encodes a protein of unknown function was only detected in our analysis but a closer visual inspection of the profiles obtained by Fraschka et al. (15) shows extensive occupancy of the gene by *Pb*HP1 (**Supplementary Figure S4B**). On the other hand, *ap2-g* and *ap2-sp3/ap2-tel* are not detected as heterochromatic in our analysis but visual inspection indicates increased *Pb*HP1 occupancy (**Supplementary Figure S4C**). These data suggest that automated detection of moderate heterochromatin enrichment remains a challenge and that conclusions about absence of epigenetic silencing in such regions should be treated with caution. Nonetheless, the two studies together reveal that *P. berghei* heterochromatin formation, maintenance and inheritance are largely hardwired throughout development.

It has been previously proposed that epigenetic reprogramming or resetting of the expression of genes involved in parasite virulence occurs during parasite passage through the vector (10, 66). However, no such evidence for a reset of virulence is detected pre- and post-meiotically in our study, which could be explained by two scenarios. Firstly, such reset is likely to start immediately after nuclear fusion as in higher eukaryotes (67) and thus may have been missed by our study design. Secondly, any such presumed reset may involve different epigenetic marks than those examined here. Finally, the virulence-reset hypothesis is based on parasites serially maintained in rodents, which are known to become more virulent. Therefore, it is probable that the observed phenotype is based on ill-managed heterochromatin maintenance undetectable by our study.

Gametocytes are poised cells that are rapidly activated and transform into respective gametes upon ingestion by a mosquito. In *P. falciparum*, heterochromatin boundaries are shown to expand outside telomeres as the parasite lifecycle progresses from ABS to gametocytes (15). However, our data do not show any major differences in heterochromatin distribution between the various *P. berghei* lifecycle stages examined. This result can be explained by the fact that genes that become heterochromatic in *P. falciparum* gametocytes and are involved iRBC remodeling (such as knob formation) have no clear orthologues in *P. berghei* and highlights differences of heterochromatin maintenance between the two parasites.

Male and female gametocytes are characterized by specific transcriptomes and proteomes (22, 68–70). However, in our analysis, differential heterochromatin occupancy between these two stages is limited to 12 genes, suggesting that epigenetic gene silencing is not a key regulatory mechanism of differential gene expression between male and female gametocytes. In addition, all these genes are located in subtelomeric regions and 11 of them belong to a multigene family, and thus some of the detected differences may be due to the method’s sensitivity. It remains to be seen whether any of the 12 genes show true expression differences between male and female gametocytes and relate to sex-specific functions.

A striking difference between male and female gametocytes is the level of H3K9ac occupancy in the 5’UTR of genes, which correlates with transcript abundance in male but not in female gametocytes. Thus it appears that gene transcription in female gametocytes is independent of H3K9ac levels. We propose that the female gametocyte is a stage of epigenetic remodeling that involves universal marking of gene promoters with H3K9ac. At the same time, genes that are involved in ribosome biogenesis and translation are downregulated, a phenomenon that together with translational mRNA repression by the DOZI-complex suggests that a multilayered mechanism regulating zygotic development operates in *Plasmodium*.

Mature oocytes in mammals, flies and worms are transcriptionally silent when passing through meiosis I preceding fertilization (71). Additionally, early embryonic development depends solely on maternally deposited proteins and RNAs and coincides with low or undetectable transcription (72). Zygotic transcription often resumes after several hours or days post fertilization, a process called maternal-to-zygotic transition accompanied by zygotic genome activation, and in flies, for example, is controlled by a master transcription factor that is not necessarily associated with H3K9ac (73). Additionally, epigenetic reprogramming occurs before and after fertilization, ensuring the transition from a highly specific cell type back to a totipotency state able to form a new organism. However, direct comparison of malaria parasite female gametocytes to oocytes of multicellular organisms is complicated by the fact that malaria parasites undergo meiosis directly after fertilization (74), in contrast to metazoan cells where meiosis precedes fertilization. Nonetheless, it is tempting to hypothesise that, similar to higher eukaryotes, (i) the female gametocyte is transcriptionally poised and (ii) transcription in the ookinete is resumed only from a selected set of genes. While there is no evidence so far that the female gametocyte is transcriptionally poised, there is evidence for the second part of the hypothesis. Firstly, when expression of a fluorescent reporter protein is controlled by an ookinete-specific gene promoter the protein is detected from both the maternal and paternal genomes in the first 24 hours after fertilization (75). Secondly, if expression of a fluorescent reporter protein is controlled by an housekeeping gene promoter the protein is detected only from the maternal genome (75). This could either mean that the paternal genome is silenced, or more likely, that both parental genomes are transcriptionally poised, and the detected expression stems from inherited maternal mRNA.

Two histone variants associated with active promoters in *P. falciparum* ABS parasites (H2A.Z and H2B.Z) (14, 76) are significantly downregulated in female gametocytes in this study (**Supplementary Table 2**). As mRNA is stored in the RNP complex in female gametocytes, we do not know which mRNA is translated into protein and which mRNA is translationally poised at this stage. Thus, it will be interesting to see if either female gametocytes are have a low occurrence of H2A.Z and H2B.Z, or if these histone variants are not deposited into the chromatin during the early development of the zygote. Proteomic studies identified both H2A.Z and H2B.Z in female gametocytes in both *P. berghei and P. falciparum* (68, 69), pointing towards the latter possibility. Supportively, in mice, H2A.Z remains undetectable in embryonic chromatin before the late 2-cell stage (77). Thus, we propose a scenario where the female gametocyte is transcriptionally poised, and where transcription of stage-specific genes is resumed in the ookinete stage.

## ACKNOWLEDGEMENTS

The authors wish to acknowledge the support from the FACS-facility from Imperial College London, namely Jess Rowley and Jane Srivastava. Andy Brockman for extracting the UTRs from the *P. berghei* ANKA genome, Tony Brooks and Paola Niola from UCL for help with RNA sequencing. Bob MacCallum, Amie Jaye and Dan Lawson for assistance in all things computational. Till S. Voss for sharing the *Pb*HP1 antibody before its publication. We are grateful to Jen Hillman for guidance/assistance on usegalaxy.org and the Freiburg Galaxy team for their assistance using usegalaxy.eu.

## Authors’ contributions

conceptualization, K.W., R.B., D.V., G.K.C.; investigation, K.W.; formal analysis, K.W., S. F.; resources, R.B, D.V., G.K.C; writing – original draft, K.W.; writing – review & editing, K.W., S.F., D.V., R.B., G.K.C.; visualization, K.W.

## Ethics approval

All animal procedures were carried out in accordance with the Animal Scientifics Procedures Act 1986 under the UK Home Office Licenses PLL70/7185 and PPL70/8788.

