## Supplemental Figures for "Epigenetic regulation underlying *Plasmodium berghei* gene expression during its developmental transition from host to vector"

### SUPPLEMENTARY DATA

#### Supplementary Figure 1. RNA sequencing of P. berghei ABS, FG, MG and OOK

(A) No GC-bias according to (42) was detected when processing genomic DNA alongside the RNA samples. Data was analysed in non-overlapping 300bp bins and the number of reads according to the underlying GC-percentage of the regions is plotted (upper boxplot). Lower panel shows read counts normalised to the expected read counts according to the underlying GC-value.

(B) Principal component analysis of aligned RNAseq data of all *P. berghei* life stages investigated including the gDNA control. Aligned and sorted reads of each sample were grouped into non-overlapping 1000bp bins. Triplicates cluster together, and transcriptionally, male gametocytes are similar to asexual blood stage parasites, whereas female gametocytes are more closely related to ookinetes.

#### Supplementary Figure 2. A unique heterochromatic dynein gene in *P. berghei*.

(A) Heat map of all 22 dynein genes and their respective H3K9ac occupancy and relative transcripts for each life cycle stage. H3K9ac enrichment for each gene locus is shown as $log_2$-transformed H3K9ac ChIP/input (1000bp upstream of ATG the ORF, and 500bp downstream of stop codon, respectively). Pink colour indicates relative transcripts (in FPKM).

(B) Bar plot of the left arm of chromosome 6. Closed box represents gene of interest, arrows indicate the orientation of each gene. Open boxes indicate neighbouring genes. Peaks correspond to log2 transformed data of either PbHP1-ChIP/input or H3K9ac-ChIP/input. $Log_2$ scale for *Pb*HP1 is -3 to 3 and -2 to 2 for H3K9ac, respectively. Numbers indicate relative transcripts (in FPKM) for each stage.

**Supplementary Figure 3. Heterochromatin changes between different life cycle stages and parasite isolates.**

Screenshots of *Pb*HP1 tracks of genes from Figure 2A that were differently heterochromatically marked in our analysis. Gene of interest is shown as a black box, and the arrow indicates its orientation. Bars above each box represent 5kb length. $Log_2$ scale for *Pb*HP1 is (-3 to 3).

**Supplementary Figure 4. Heterochromatin distribution in asexual blood stages is very similar to the *Pb*ANKA isolate used in Fraschka et al. (2018).**

(A) Chrosomosomal location of each heterochromatic gene from this study and the study from Fraschka and colleagues. Each filled dot represents a gene. Colour code shows if a gene was found to be heterochromatic in both studies (blue), this study (green) or the study from Fraschka et al (red). Taken both studies together, only four heterochromatic genes are not located at subtelomeric regions (highlighted with an arrow).

(B) and (C) Bar chart of or two of the genes highlighted in A. Small arrows underneath the gene indicates direction of transcription. $Log_2$ scale for *Pb*HP1 is (-3 to 3).

**Supplementary Figure 5. H3K9ac occupancy in female gametocytes in different gene sets.**

(A) Heatmap showing H3K9ac occupancy of the 5'UTR of translationally repressed genes (DOZI-controlled) (65) compared to all euchromatic genes.

(B) Box plot showing mean 5'UTR values from the two gene groups from (A). DOZI-controlled genes have significantly less H3K9ac coverage in their 5'UTR. Asterisks mark significance (Mann-Whitney test, p =0.0004).

(C) Heatmap showing H3K9ac occupancy of the 5'UTR of AP2-O-controlled genes (23) compared to all euchromatic genes.

(D) Box plot showing mean 5'UTR values from the two gene groups from (C). No statistically significant difference has been found (Mann-Whitney test (ns), p=0.6245).

**Supplementary Table 1. PbHP1 enrichment and relative transcripts of heterochromatin genes.**

Mean PbHP1 values for all genes found to be enriched by PbHP1. FPKM (mean of three biological replicates) is shown.

**Supplementary Table 2. List of genes that are significantly up- and downregulated in FG compared to all other stages.**

**Supplementary Table 3. Go-terms associated with up- or downregulated genes in female gametocytes.**

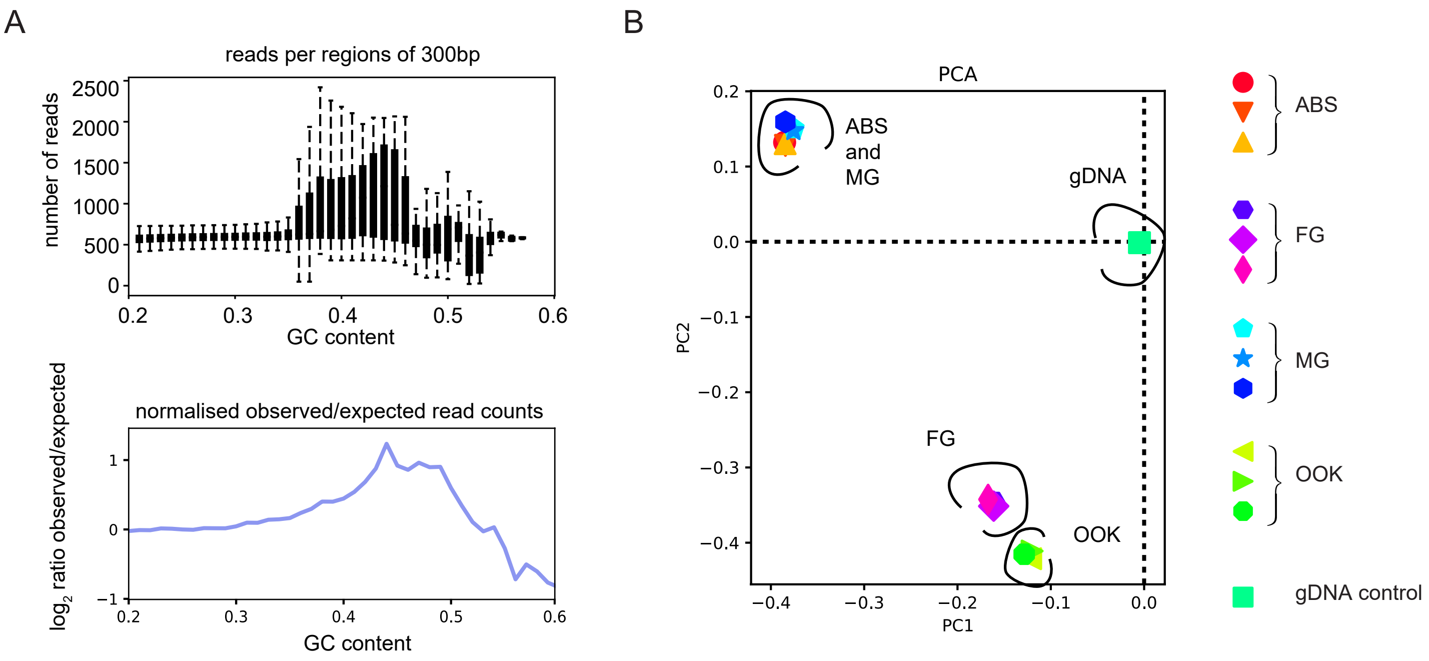


Supplementary Figure 1

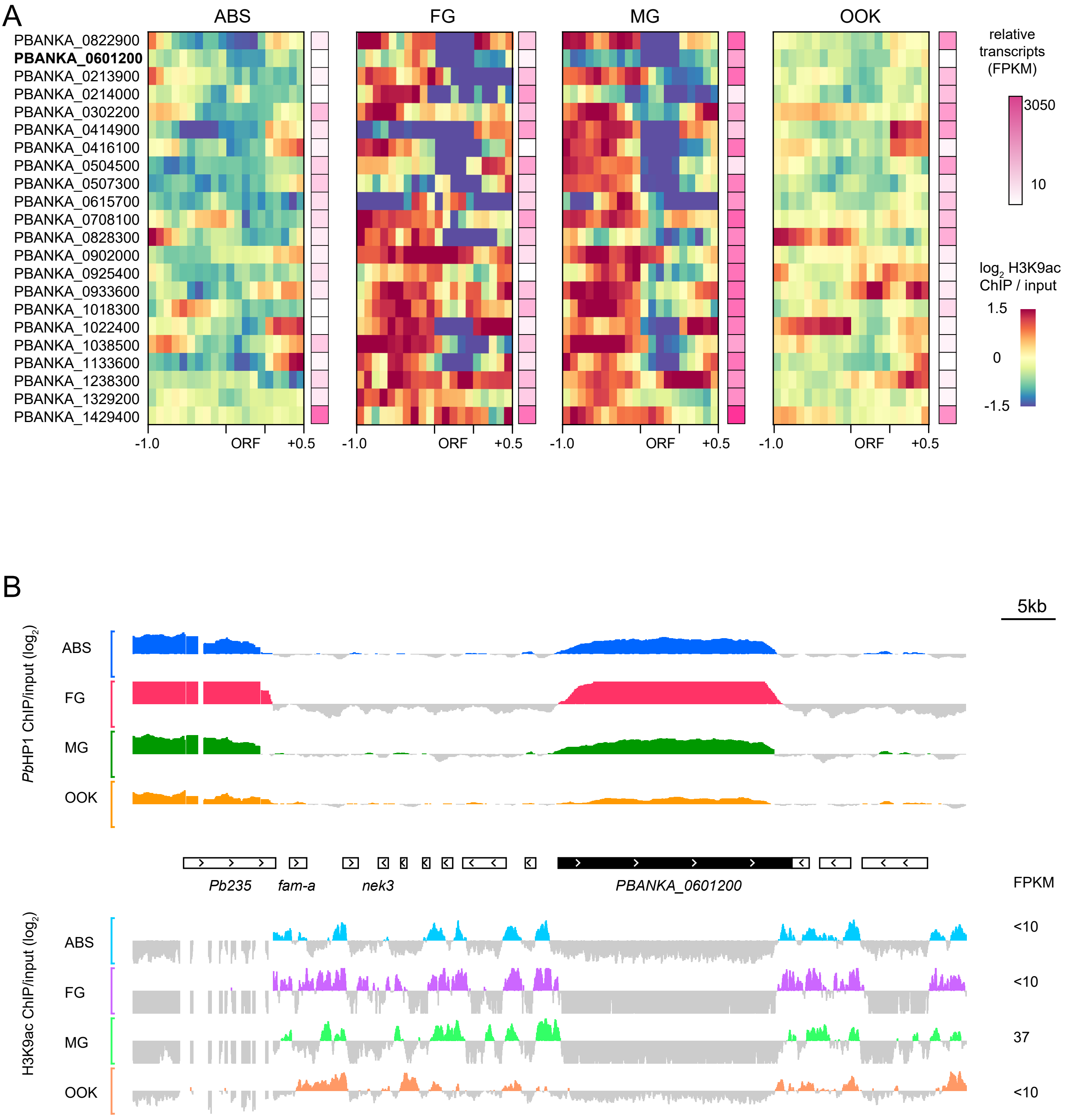


Supplementary Figure 2

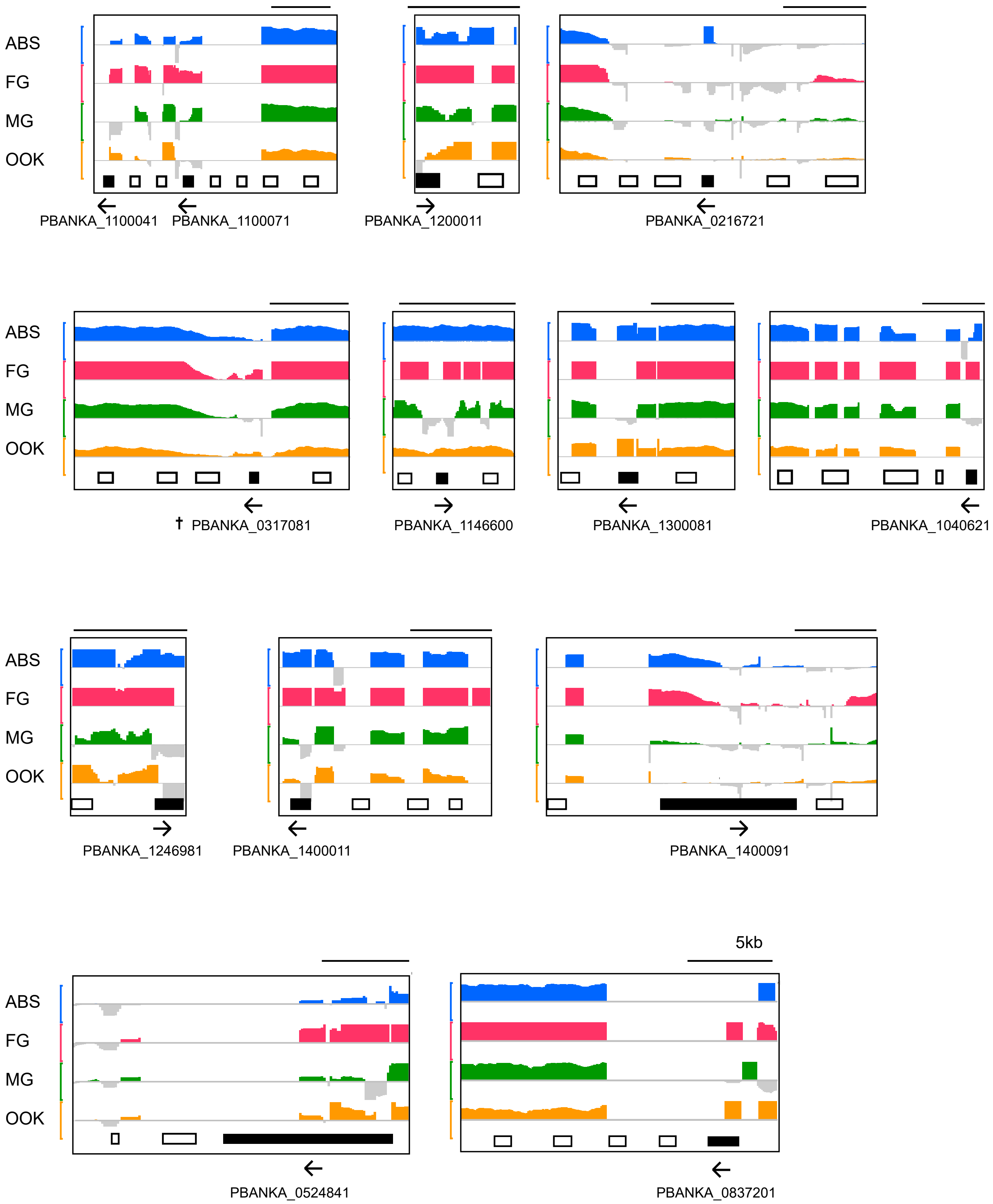


Supplementary Figure 3

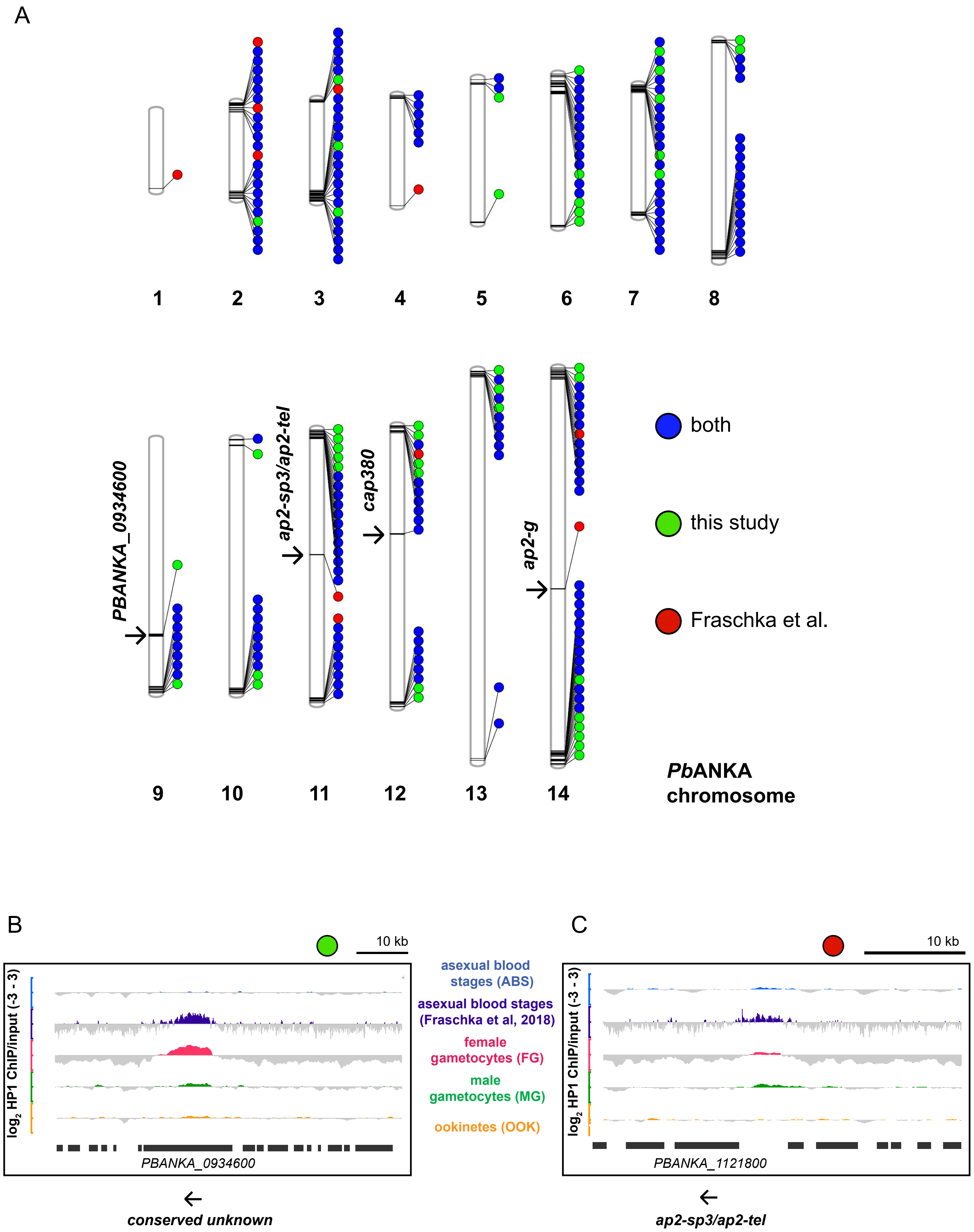


Supplementary Figure 4

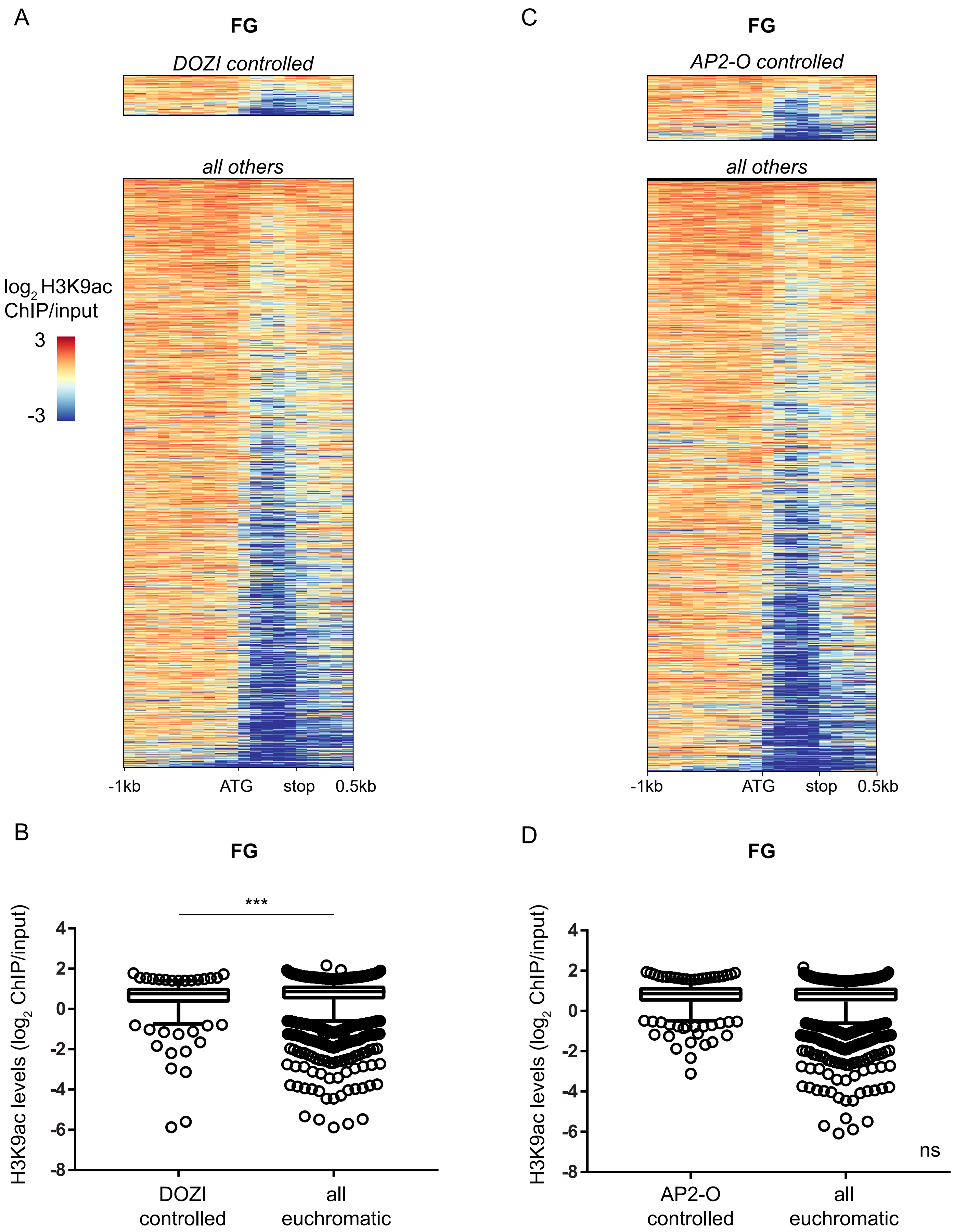


Supplementary Figure 5
